# Harmonization of whole genome sequencing for outbreak surveillance of Enterobacteriaceae and Enterococci

**DOI:** 10.1101/2020.11.20.392399

**Authors:** Casper Jamin, Sien de Koster, Stefanie van Koeveringe, Dieter de Coninck, Klaas Mensaert, Katrien de Bruyne, Natascha Perales Selva, Christine Lammens, Herman Goossens, Christian Hoebe, Paul Savelkoul, Lieke van Alphen, and the i-4-1-Health Study Group

**Author notes:** Corresponding Author: Casper Jamin PhD candidate, Department of Medical Microbiology, Care and Public Health Research Institute (CAPHRI), Maastricht University Medical Center+, Maastricht, The Netherlands. **i-4-1-Health Study Group** Lieke van Alphen (Maastricht University Medical Center+, Maastricht, the Netherlands), Nicole van den Braak (Avans University of Applied Sciences, Breda, the Netherlands), Caroline Broucke (Agency for Care and Health, Brussels, Belgium), Anton Buiting (Elisabeth-TweeSteden Hospital, Tilburg, the Netherlands), Liselotte Coorevits (Ghent University Hospital, Ghent, Belgium), Sara Dequeker (Agency for Care and Health, Brussels, Belgium and Sciensano, Brussels, Belgium), Jeroen Dewulf (Ghent University, Ghent, Belgium), Wouter Dhaeze (Agency for Care and Health, Brussels, Belgium), Bram Diederen (ZorgSaam Hospital, Terneuzen, the Netherlands), Helen Ewalts (Regional Public Health Service Hart voor Brabant, Tilburg, the Netherlands), Herman Goossens (University of Antwerp, Antwerpen, Belgium and Antwerp University Hospital, Antwerp, Belgium), Inge Gyssens (Hasselt University, Hasselt, Belgium), Casper den Heijer (Regional Public Health Service Zuid-Limburg, Heerlen, the Netherlands), Christian Hoebe (Maastricht University Medical Center+, Maastricht, the Netherlands and Regional Public Health Service Zuid-Limburg, Heerlen, the Netherlands), Casper Jamin (Maastricht University Medical Center+, Maastricht, the Netherlands), Patricia Jansingh (Regional Public Health Service Limburg Noord, Venlo, the Netherlands), Jan Kluytmans (Amphia Hospital, Breda, the Netherlands and University Medical Center Utrecht, Utrecht University, Utrecht, the Netherlands), Marjolein Kluytmans-van den Bergh (Amphia Hospital, Breda, the Netherlands and University Medical Center Utrecht, Utrecht University, Utrecht, the Netherlands), Stefanie van Koeveringe (Antwerp University Hospital, Antwerp, Belgium), Sien De Koster (University of Antwerp, Antwerp, Belgium), Christine Lammens (University of Antwerp, Antwerp, Belgium), Isabel Leroux-Roels (Ghent University Hospital, Ghent, Belgium), Hanna Masson (Agency for Care and Health, Brussel, Belgium), Ellen Nieuwkoop (Elisabeth-TweeSteden Hospital, Tilburg, the Netherlands), Anita van Oosten (Admiraal De Ruyter Hospital, Goes, the Netherlands), Natascha Perales Selva (Antwerp University Hospital, Antwerp, Belgium), Merel Postma (Ghent University, Ghent, Belgium), Stijn Raven (Regional Public Health Service West-Brabant, Breda, the Netherlands), Veroniek Saegeman (University Hospitals Leuven, Leuven, Belgium), Paul Savelkoul (Maastricht University Medical Center+, Maastricht, the Netherlands), Annette Schuermans (University Hospitals Leuven, Leuven, Belgium), Nathalie Sleeckx (Experimental Poultry Centre, Geel, Belgium), Arjan Stegeman (Utrecht University, Utrecht, the Netherlands), Tijs Tobias (Utrecht University, Utrecht, the Netherlands), Paulien Tolsma (Regional Public Health Service Brabant Zuid-Oost, Eindhoven, the Netherlands), Jacobien Veenemans (Admiraal De Ruyter Hospital, Goes, the Netherlands), Dewi van der Vegt (PAMM Laboratory for Pathology and Medical Microbiology, Veldhoven, the Netherlands), Martine Verelst (University Hospitals Leuven, Leuven, Belgium), Carlo Verhulst (Amphia Hospital, Breda, the Netherlands), Pascal De Waegemaeker (Ghent University Hospital, Ghent, Belgium), Veronica Weterings (Amphia Hospital, Breda, the Netherlands), Clementine Wijkmans (Regional Public Health Service Hart voor Brabant, Tilburg, the Netherlands), Patricia Willemse-Smits (Elkerliek Hospital, Helmond, the Netherlands), Ina Willemsen (Amphia Hospital, Breda, the Netherlands).

## Abstract

Whole genome sequencing (WGS), is becoming the facto standard for bacterial typing and outbreak surveillance of resistant bacterial pathogens. We performed a three-center ring trial to assert if inter-laboratory harmonization of WGS is achievable, for this goal. To this end, a set of 30 bacterial isolates comprising of various species belonging to the Enterobacteriaceae and Enterococcus genera were selected and sequenced using the same protocol on the Illumina MiSeq platform in each individual centre. All generated sequencing data was analysed by 1 centre using BioNumerics (6.7.3) for i) genotyping origin of replications & antimicrobial resistance genes, ii) core-genome (cgMLST) for *E. coli* and *K. pneumoniae* & whole-genome multi locus sequencing typing (wgMLST) for all species. Additionally, a split k-mer analysis was performed to determine the number of SNPs between samples. A precision of 99.0% and an accuracy of 99.2% was achieved for genotyping. Based on cgMLST, only in 2/27 and 3/15 comparisons a discrepant allele was called between two genomes, for *E. coli* and *K. pneumonia,* respectively. Based on wgMLST, the number of discrepant alleles ranged from 0 to 7 (average 1.6). For SNPs, this ranged from 0-11 SNPs (average 3.4). Furthermore, we demonstrate that using different *de novo* assemblers to analyse the same dataset introduces up to 150 SNPs, which surpasses most thresholds for bacterial outbreaks. This shows the importance of harmonisation of data processing surveillance of bacterial outbreaks. Summarizing, multi-center WGS for bacterial surveillance is achievable, but only if protocols are harmonized.

## Introduction/background

The surge and dissemination of antimicrobial resistance (AMR) has grown to an issue of worldwide proportions. Key objectives in fighting this problem are preventing the spread of AMR bacteria and improving surveillance of these resistant microbes. The prevention of (hospital-acquired) infections with resistant microbes^1^ is important to reduce the spread of antimicrobial resistant micro-organisms. Routine surveillance by molecular typing can aid in this prevention and thus in the fight against AMR, as outlined by the global action plan of the World Health Organization. ESKAPE pathogens (*Enterococcus faecium, Staphylococcus aureus, Klebsiella pneumoniae, Acinetobacter baumannii, Pseudomonas aeruginosa and Enterobacter sp*.) are of major interest as they are the leading cause of hospital-related infections and outbreaks. Furthermore, reports show that the number of infections by resistant microorganisms have been on the rise in recent years. Infections by multi-drug resistant (MDR) bacteria are associated with an increase in economic burden^2^ and negative patient outcomes such as morbidity and mortality^3,4^.

To determine the spread of resistance and resistant microbes, different molecular typing methods are being applied. Older, established typing methods for outbreak surveillance, such as PFGE, AFLP, MLST and MLVA are slowly being replaced by whole genome sequencing (WGS). The introduction of WGS to the field of bacterial typing and spread of AMR has set a new standard for discriminatory power and accuracy, as it encompasses a comprehensive view of the bacterial core and accessory genome. This gives rise to the possibility to determine clonal relatedness in a more discriminatory fashion, and at the same time provide data on resistance genes, plasmids and virulence-potential, which would otherwise require a combination of other methods^5–8^. Current methods to determine phylogeny are based on core/whole genome multi-locus sequence typing (cgMLST, wgMLST)^9,10^ or single nucleotide polymorphisms (SNPs)^11–13^.

Approaches like cgMLST and wgMLST determine the phylogeny among bacterial strains based on difference in allelic profile in either the core genome or the entire genome, respectively. All coding sequences (CDS) or loci are identified using tools such as Prodigal^14^. Then, all variants of each locus are assigned a unique allele number and the complete set of allele numbers is called the allelic profile. The genetic distance is calculated by counting the number of discrepant alleles between two strains. A relative genetic distance can also be calculated by dividing the number of discrepant alleles by the number of alleles that were compared. Next to commercial packages for cgMLST and wgMLST analyses, such as BioNumerics or SeqSphere, open source options are available as well, like ChewBBACA^10^ and Enterobase^15^.

Inferring phylogeny based on SNPs can be performed by three different methods. i) Alignment to a reference genome (Snippy ^11^). ii) (core-) genome alignment (MAUVE^16^ or Harvest Suite^17^). iii) alignment-free methods based on using the entire collection of subsequences of a sequence of length k: k-mer(kSNP^18^ or SKA^13^).

Currently, only few studies have described clonal cluster thresholds definitions using cgMLST, wgMLST or SNP-based methods. Generally, these studies determine the thresholds based on either i) previous or ongoing bacterial outbreaks in hospitals and in the food production chain, or ii) by means of follow-up on human carriers of these pathogens over time. Furthermore, most of these studies only describe single clone outbreaks, which can hamper the interpretation when these thresholds are applied to different lineages of a specific species, as some clinically relevant lineages might be more clonal than others, requiring different thresholds. One of the first reports on the use of WGS for bacterial outbreak analysis was on methicillin-resistant *Staphylococcus aureus* (MRSA) in 2013 in a neonatal intensive care unit. Next to standard assessment of epidemiological data and antibiograms, WGS was performed to resolve this putative outbreak^19^. In that study, a maximum of 20 SNPs was observed among the MRSA strains found in the outbreak. For the foodborne pathogen *E. coli* O157:H7, the Public health agency Canada evaluated WGS for outbreak detection^20^. Here, 250 strains, comprising eight different outbreaks that were previously typed by older methods were retrospectively sequenced and analyzed using wgMLST and SNP analyses. These two methods inferring phylogenetic relations among strains were reported to be in excellent concordance with each other. Additionally, they reported that all strains per outbreak fell within a cutoff of 5 SNPs or 10 allele differences (on wgMLST basis). Schürch *et al.* suggested in their review various clonal cluster thresholds based on wgMLST or SNP analyses for a few common bacterial pathogens in outbreak situations^9^.

Kluytmans-van den Bergh *et al.* recently determined clonal-cutoffs based on cgMLST and wgMLST for four extended-spectrum beta-lactamase-producing Enterobacteriaceae (ESBL-E)*: E. coli, K. pneumoniae*, *Citrobacter ssp*. and *Enterobacter cloacae* complex^21^. Here, strains were classified as epidemiologically linked when these were cultured from a single patient in a 30-day time window and when they belonged to the same seven-gene sequence type. Subsequently, the genetic distance (here defined as number of discrepant alleles divided by the number of alleles compared) was compared among all strains, and clonal thresholds were determined by the lowest genetic distance possible that included all epidemiologically linked strains.

The goal of the i-4-1-Health study is to assess the prevalence and spread of resistant bacteria among humans and animals in the Dutch-Belgian border^22^. Within a one-year period, patients in hospitals and in long-term healthcare facilities, infants at day-care facilities, as well as broilers and weaned pigs were screened for gut or rectal carriage of ESBL-producing, ciprofloxacin-resistant or carbapenemase-producing Enterobacteriaceae and vancomycin-resistant *Enterococci*. This one-health approach could provide insights in the prevalence and spread of resistant bacteria between and within these separate domains. In this project, WGS data was generated in three independent locations, and thus, inter-laboratory reproducibility needed to be assed to allow the comparison of this data. To standardize the WGS results and interpretation, efforts were made to harmonize the WGS protocols, both for the wet-lab procedures, as well as the bioinformatics analysis.

Here, we harmonized the inter-laboratory reproducibility of WGS for outbreak surveillance and genotyping of AMR and origin of replication (ORI) of plasmids for a selection of AMR bacteria frequently encountered in hospital-related infections and AMR surveillance within the I-4-1-Health project. As the implementation of WGS for routine outbreak surveillance is particularly dependent on standardized methodology, we evaluated the technical variation in phylogenetic comparison using a commercially available wgMLST tool in BioNumerics and an open source reference free SNP-based tool called SKA^13^.

## Materials and Methods

### Selection of strains

In total 30 resistant bacterial strains were selected based on their extended-spectrum bèta-lactamase (ESBL) or carbapenemase activity, or based on ciprofloxacin or vancomycin resistance phenotype. The complete collection of strains consisted of 9 *Escherichia coli*; 5 *Klebsiella pneumonia*; 4 *Citrobacter subspecies*; 4 *Enterobacter cloacae;* 2 *Klebsiella oxytoca;* 2 *Enterobacter aerogenes;* 2 *Enterococcus faecalis* and 2 *Enterococcus faecium*. Six strains (2 *Escherichia coli,* 2 *Klebsiella pneumoniae* and 2 *Enterobacter cloacae*) were previously collected^21^ and were kindly provided by the SoM study-group, and 20 strains during the i-4-1-Health study^22^. The *E. faecium* and *E. faecalis* strains were from a previously acquired collection, stored at Antwerp University. The strains were collected from perianal swabs of hospitalized patients (21) and clients in nursing homes (6), and from feces from broilers (2) and weaned pigs (1) by selective culturing. The culturing methods were described elsewhere^21,23^. All strains originated from The Netherlands and Belgium. An overview of strains and their origin is available in table 1. Strains were inoculated from −80°C on Mueller Hinton II agar (BD, Franklin Lakes, NJ, USA) and sent to the participating institutes. The 30 strains were divided in three sets of ten strains. Each set was sequenced once by each center with a six-month interval between each set.

**Table 1.**
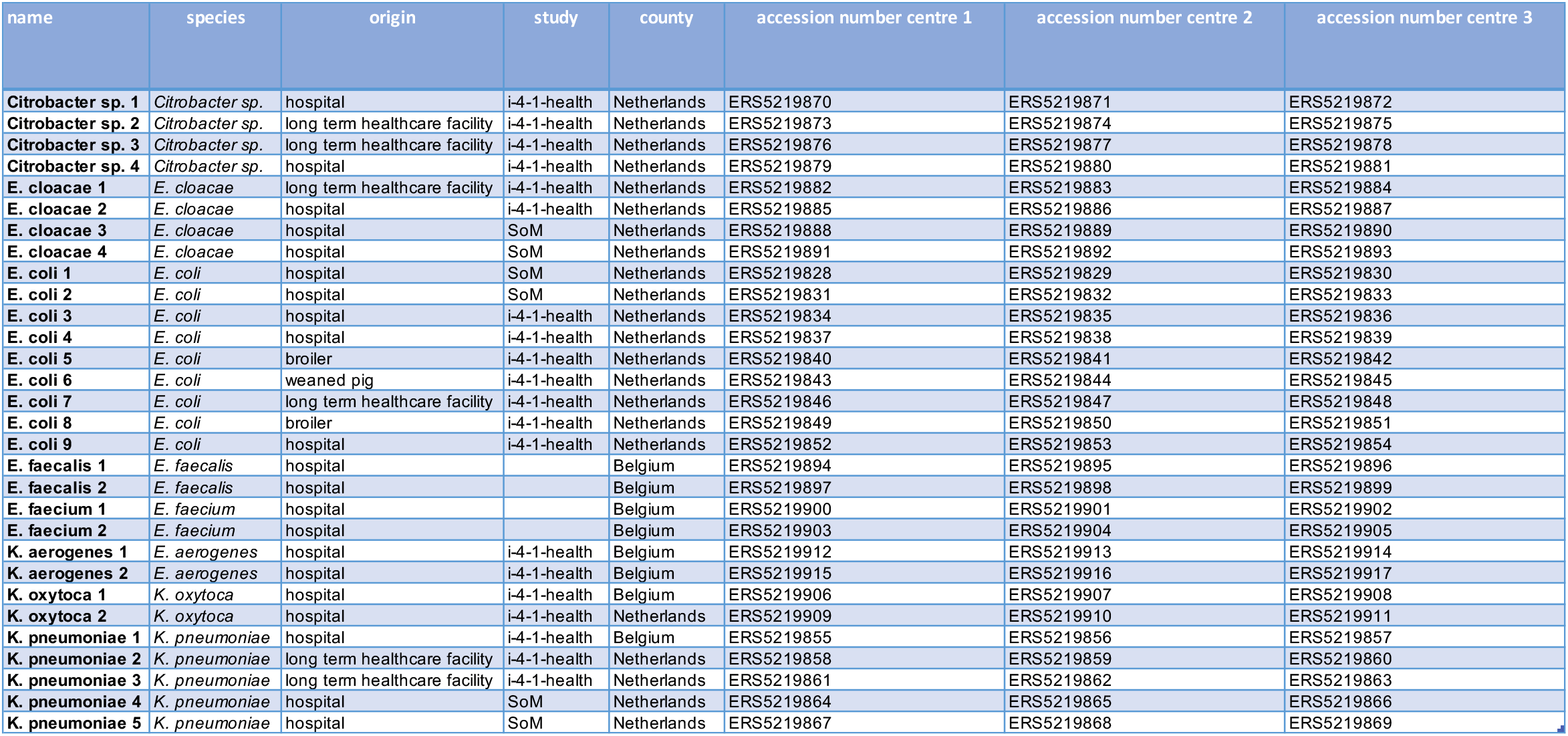
Overview of strains used in this study.

| name | species | origin | study | county | accession number centre 1 | accession number centre 2 | accession number centre 3 |
| --- | --- | --- | --- | --- | --- | --- | --- |
| **Citrobacter sp. 1** | *Citrobacter sp.* | hospital | i-4-1-health | Netherlands | ERS5219870 | ERS5219871 | ERS5219872 |
| **Citrobacter sp. 2** | *Citrobacter sp.* | long term healthcare facility | i-4-1-health | Netherlands | ERS5219873 | ERS5219874 | ERS5219875 |
| **Citrobacter sp. 3** | *Citrobacter sp.* | long term healthcare facility | i-4-1-health | Netherlands | ERS5219876 | ERS5219877 | ERS5219878 |
| **Citrobacter sp. 4** | *Citrobacter sp.* | hospital | i-4-1-health | Netherlands | ERS5219879 | ERS5219880 | ERS5219881 |
| **E. cloacae 1** | *E. cloacae* | long term healthcare facility | i-4-1-health | Netherlands | ERS5219882 | ERS5219883 | ERS5219884 |
| **E. cloacae 2** | *E. cloacae* | hospital | i-4-1-health | Netherlands | ERS5219885 | ERS5219886 | ERS5219887 |
| **E. cloacae 3** | *E. cloacae* | hospital | SoM | Netherlands | ERS5219888 | ERS5219889 | ERS5219890 |
| **E. cloacae 4** | *E. cloacae* | hospital | SoM | Netherlands | ERS5219891 | ERS5219892 | ERS5219893 |
| **E. coli 1** | *E. coli* | hospital | SoM | Netherlands | ERS5219828 | ERS5219829 | ERS5219830 |
| **E. coli 2** | *E. coli* | hospital | SoM | Netherlands | ERS5219831 | ERS5219832 | ERS5219833 |
| **E. coli 3** | *E. coli* | hospital | i-4-1-health | Netherlands | ERS5219834 | ERS5219835 | ERS5219836 |
| **E. coli 4** | *E. coli* | hospital | i-4-1-health | Netherlands | ERS5219837 | ERS5219838 | ERS5219839 |
| **E. coli 5** | *E. coli* | broiler | i-4-1-health | Netherlands | ERS5219840 | ERS5219841 | ERS5219842 |
| **E. coli 6** | *E. coli* | weaned pig | i-4-1-health | Netherlands | ERS5219843 | ERS5219844 | ERS5219845 |
| **E. coli 7** | *E. coli* | long term healthcare facility | i-4-1-health | Netherlands | ERS5219846 | ERS5219847 | ERS5219848 |
| **E. coli 8** | *E. coli* | broiler | i-4-1-health | Netherlands | ERS5219849 | ERS5219850 | ERS5219851 |
| **E. coli 9** | *E. coli* | hospital | i-4-1-health | Netherlands | ERS5219852 | ERS5219853 | ERS5219854 |
| **E. faecalis 1** | *E. faecalis* | hospital |  | Belgium | ERS5219894 | ERS5219895 | ERS5219896 |
| **E. faecalis 2** | *E. faecalis* | hospital |  | Belgium | ERS5219897 | ERS5219898 | ERS5219899 |
| **E. faecium 1** | *E. faecium* | hospital |  | Belgium | ERS5219900 | ERS5219901 | ERS5219902 |
| **E. faecium 2** | *E. faecium* | hospital |  | Belgium | ERS5219903 | ERS5219904 | ERS5219905 |
| **K. aerogenes 1** | *E. aerogenes* | hospital | i-4-1-health | Belgium | ERS5219912 | ERS5219913 | ERS5219914 |
| **K. aerogenes 2** | *E. aerogenes* | hospital | i-4-1-health | Belgium | ERS5219915 | ERS5219916 | ERS5219917 |
| **K. oxytoca 1** | *K. oxytoca* | hospital | i-4-1-health | Belgium | ERS5219906 | ERS5219907 | ERS5219908 |
| **K. oxytoca 2** | *K. oxytoca* | hospital | i-4-1-health | Netherlands | ERS5219909 | ERS5219910 | ERS5219911 |
| **K. pneumoniae 1** | *K. pneumoniae* | hospital | i-4-1-health | Belgium | ERS5219855 | ERS5219856 | ERS5219857 |
| **K. pneumoniae 2** | *K. pneumoniae* | long term healthcare facility | i-4-1-health | Netherlands | ERS5219858 | ERS5219859 | ERS5219860 |
| **K. pneumoniae 3** | *K. pneumoniae* | long term healthcare facility | i-4-1-health | Netherlands | ERS5219861 | ERS5219862 | ERS5219863 |
| **K. pneumoniae 4** | *K. pneumoniae* | hospital | SoM | Netherlands | ERS5219864 | ERS5219865 | ERS5219866 |
| **K. pneumoniae 5** | *K. pneumoniae* | hospital | SoM | Netherlands | ERS5219867 | ERS5219868 | ERS5219869 |

### DNA isolation and Whole genome sequencing

The DNA isolation procedure and WGS procedure was performed as follows: DNA was extracted using the MasterPure DNA isolation kit (Lucigen) or MasterPure Gram Positive DNA purification kit (Lucigen). Sequencing libraries were prepared using NexteraXT (Illumina), freely choosing index primers. Libraries were sequenced on the Illumina MiSeq platform in paired end 2×250 base pairs (bp) reads using MiSeq V2 cartridge. Preferably, each set of strains was subjected to WGS in a single run. Acceptance criteria for WGS were a *de novo* assembly with an average coverage higher than 30 and less than 1000 contigs, as reported in BioNumerics (7.6.3). Samples not fulfilling acceptance criteria were re-sequenced. The accession numbers for the raw sequencing data are available in table 1. Generated data was shared among institutes and analysis of all generated datasets (n=90) was performed in one institute.

### Statistical analysis

Statistical analyses were done using scipy.stats module (V1.3.1)^24^ and the statsmodel.api package in Python (v3.7).

### cgMLST and wgMLST allele calling and genotyping

Raw sequencing reads were assembled using a custom pipeline in BioNumerics (7.6.3) employing SPAdes^25^ (v3.7.0) for its *de novo* assembly. From the raw reads and the *de novo* assembly, alleles were called for core genome and whole genome MLST (cgMLST/wgMLST). In BioNumerics, cgMLST schemes were only available for *E. coli* and *K. pneumoniae* consisting of 2513 and 634 fixed loci, respectively. Pairwise allelic distance was determined by counting the number of discrepant allele variants between two datasets, ignoring loci that were not present in both datasets. Resistance genes and Origins of replication (ORI) were determined using BLAST^26^ and two custom databases based on Resfinder^27^ and PlasmidFinder^28^. AMR genes were called with a using 90% identity and 60% length cutoff. ORIs were called using 95% identity and 60% length cutoff. In total, 90 WGS datasets were generated. As no gold standard with regard to true genotype of each strain was available, the following rules were applied: (i) If either two or three out of three datasets of a strain had a specific genotype, this was considered as a true positive observation; (ii) If only one out of three datasets of a strain had a specific genotype, this was considered as a false positive observation; (iii) If a different allelic variant was observed (i.e two blaTEM-1B and one blaTEM-116) this was noted as a discrepancy and counted as a false positive.

### wgSNP analysis

To determine the best *de novo* assembler to use for wgSNP analysis, the assembler generating the least amount of pairwise SNPs (using SKA), among assemblies of the same strain, was chosen. To avoid complexity, only the *E. coli* dataset of this study was used. The following assemblers were used: I) SPAdes (v3.14.0)^25^, II) SKESA (parameters: ‘—use-paired_end’, v2.3.0)^29^, III) Megahit^30^ (v1.2.9). All tools were used in default settings, unless specified otherwise. Additionally, the assembly free method to determine SNPs straight from the raw reads, using “SKA fastq”, was also used in this comparison. The complete workflow is available at “https://github.com/MUMC-MEDMIC/assemblercompare” (v1.0). SKA^13^ was used to determine SNPs on a whole genome level, using a split k-mer length of 31. In short, pairwise SNPs were determined by generating a profile of split k-mers, in which the middle base may vary (“SKA fasta” for assembly-or “SKA fastq” for read based SNP profiling). The number of SNPs, between two datasets, was determined by comparing the split k-mer files (“SKA distance”). All data preprocessing for the SNP based data analysis was performed using Snakemake^31^ as workflow manager.

### Data availability

All raw sequencing data was deposited at EBI-ENA under BioProject PRJEB40571.

## Results

The assembly coverage, or the depth of coverage, of all strains ranged from 30 to 203 (figure 1A). The N50 score, indicative for how fragmented a de novo assembly is, ranged from 33.712bp (*E. faecium*) to 942.715 bp (*K. pneumoniae*) and showed clear species dependence (FIG 1.B). Assemblies of *E. coli, E. faecalis* and *E. faecium* showed a lower N50 score, indicating the difficulties of assembling such genomes (Figure 1B). The number of contigs also varied per species, and overall had a significant negative correlation with the sequencing depth (P < 0.01, Spearman rank correlation, FIG1.F). The number of wgMLST alleles called ranged from 1933 (*Citrobacter sp.)* alleles to 5493 (*K. pneumoniae*, Figure 1C). Furthermore, the average number of alleles per kilobases (kb) ranged from 0.41 to 0.98. A significant positive trend for the normalized allele count and sequencing depth was observed (P < 0.05, Spearman rank correlation FIG1.G). Surprisingly, mainly the *Citrobacter* datasets seemed to showed a low coding density (range 0.41 to 0.65) compared to the median of the entire dataset (0.83). Further inspection of the *Citrobacter* genome assemblies using BLAST webservice (https://blast.ncbi.nlm.nih.gov/Blast.cgi, accessed 1-4-2020), showed low homology (~85% DNA identity score) to known *Citrobacter* strains available in the NCBI database (accessed on 1-4-2020, data not shown).

**Figure 1.**
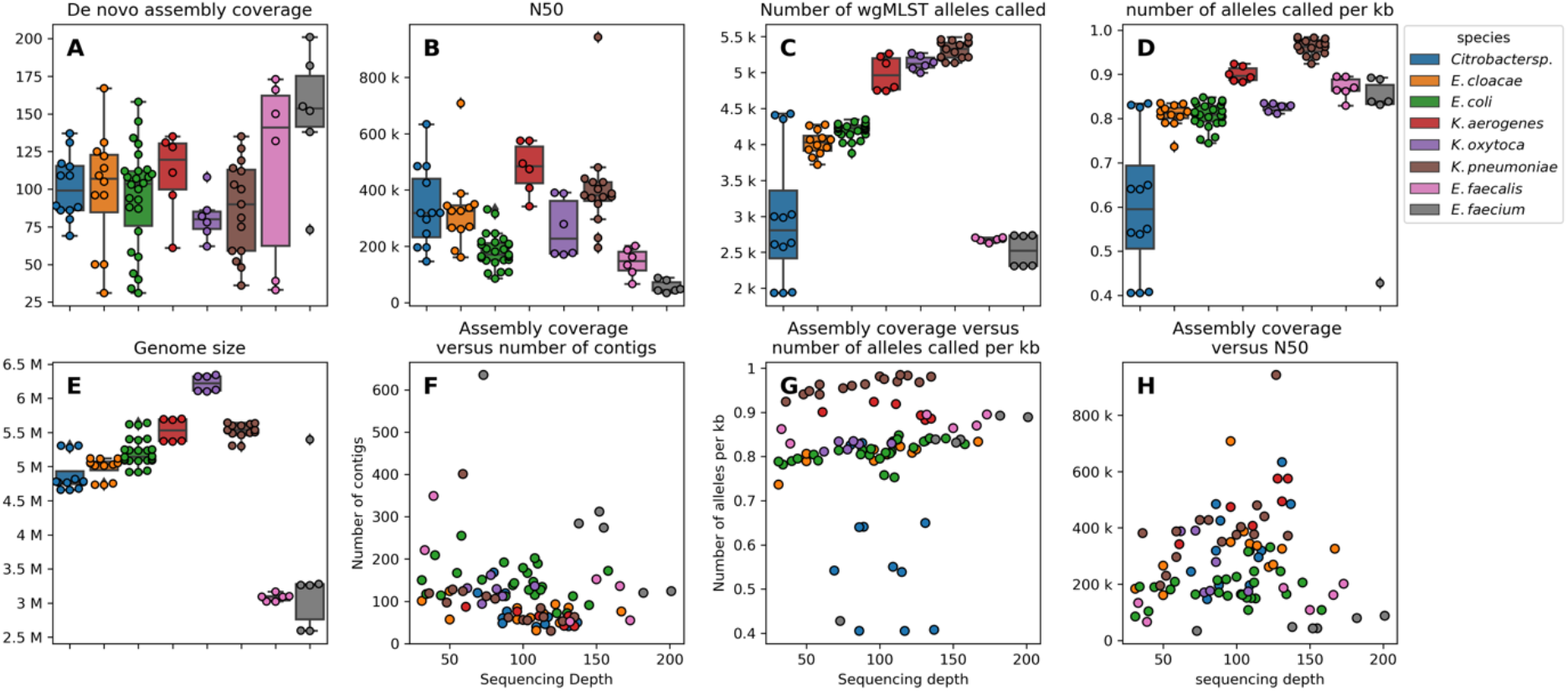
Distribution of various quality parameters pre- and post de novo assembly. Subplot A to E show boxplots with interquartile (IQ) range. Whiskers range up to 1.5 times the IQ range. All single datapoints are represented as single dots. Subplot F to G show scatterplots of relations between two quality metrics.

One dataset of *E. faecium-1* had an unusual large genome size of 5.4Mb (Figure 1E). This strain also had a higher number of contigs; (636, median of 274 for *E. faecium,* Figure 1F), and showed a lower number of alleles per kb (0.43, median of 0.84. Figure 1D) compared to the other *E. faecium* datasets. This indicates contamination in the NGS dataset of a non-*E. faecium* microbe. Manual inspection of the assembly, using BLAST webservice, showed the presence of contigs belonging to *Cutibacterium* (formerly known as *Propionibacterium*), a skin commensal and previously described as a common contaminant of NGS datasets^32–34^.

### Resistance genes and plasmid ORIs

Overall, a good consensus was obtained for the genotyping of plasmid ORIs and AMR genes (Figure 2A and 2B). A total of 973 AMR genes and ORIs were called with a precision of 99.0% and sensitivity of 99.2%. For four strains, a genotype was not called in one of the datasets. The missed genotypes were for *E. cloacae-2* a *sul1* gene, for *E. coli-6* a *tet(A)* gene, for *E. faecalis-*1 an *aac(6’)-aph(2”)* gene, and for *E. faecium-2* an *aph(2’’)-Ia* gene. For *Citrobacter-2* and *K. oxytoca-2,* a false discovery of a *blaTEM-116* was observed, as this genotype was not called in either of the other two datasets of these strains. For four strains, a discrepant genotype was called. These discrepancies were observed for *K. aerogenes-2* (*blaTEM*), for *E. cloacae-2* (*aadA*), and for *K. oxytoca-1* (*blaOXY* and *blaTEM*). Twice, an unexpected ColpVC was found in a *K. oxytoca-2* and *K. pneumoniae-4* dataset, which were from two different centers, indicating either loss of this plasmid in the other isolates of this strain, or contamination during DNA isolation or library preparation (Figure 2A).

**Figure 2A.**
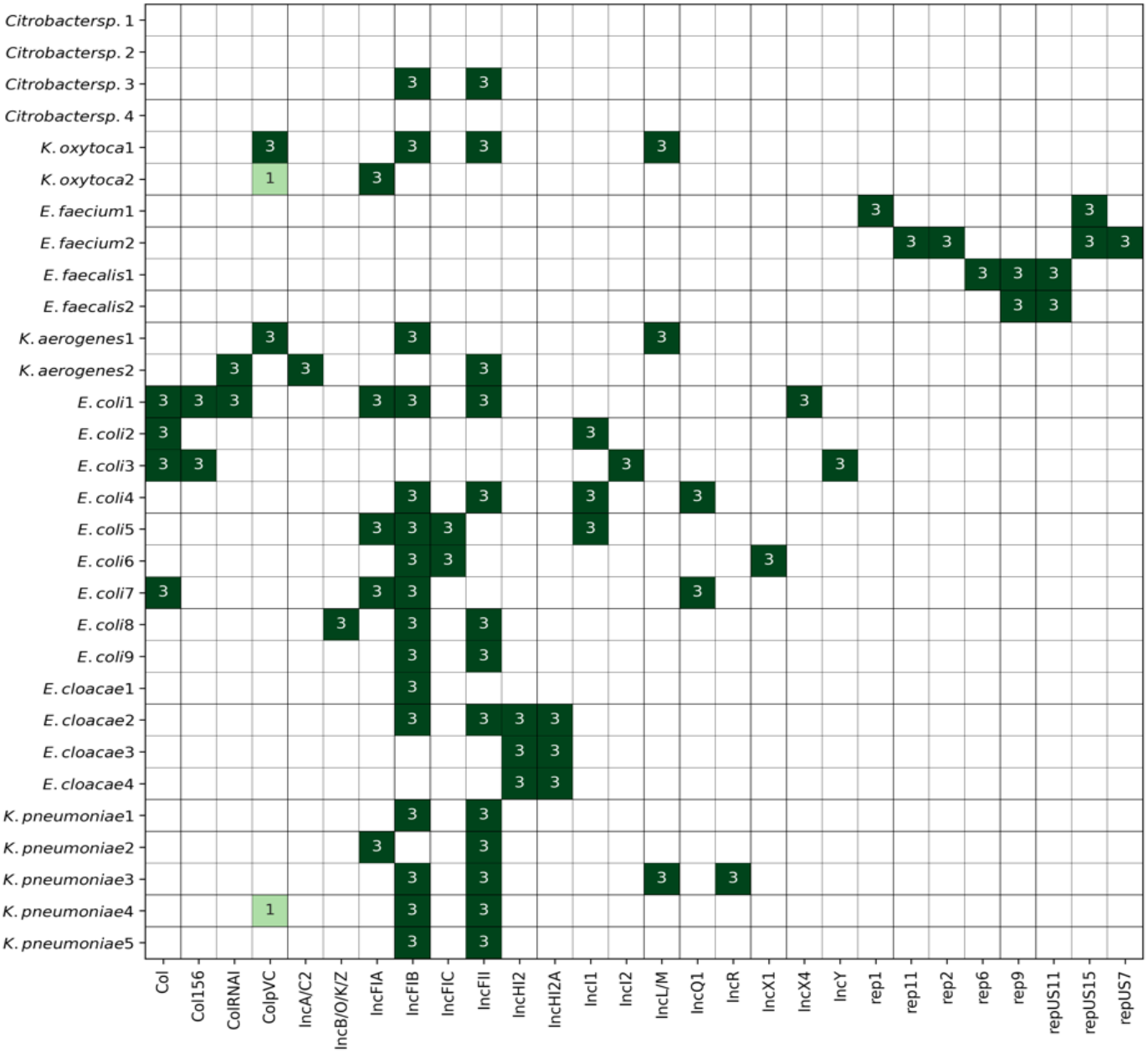
Heatmap of the number of genotype calls for various origins of replication among the isolates subjected to WGS. Genotype calls per locus was summed up for each center’s isolate if this locus was detected in their dataset.

**Figure 2B.**
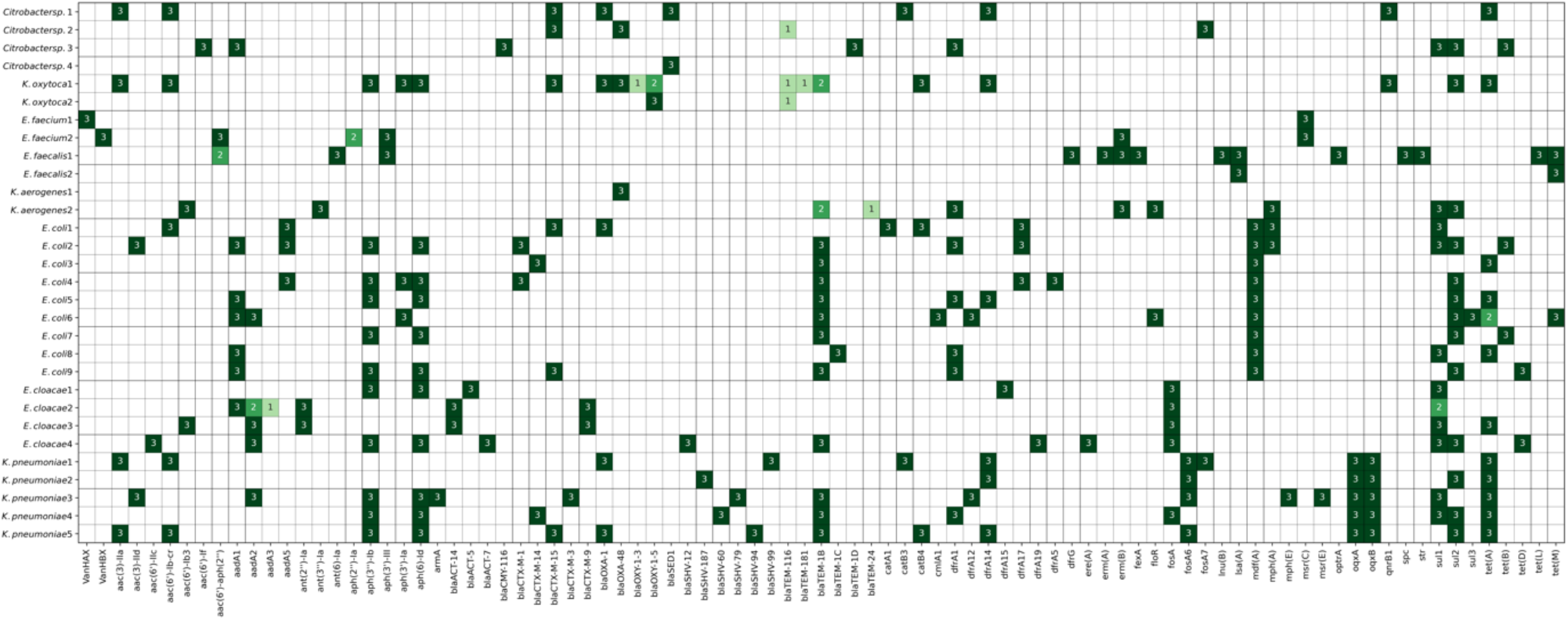
Heatmap of the number of genotype calls for various AMR genes, among the isolates subjected to WGS. Genotype calls per locus was summed up for each center’s isolate if this locus was detected in their dataset.

### Inter-laboratory variation in cgMLST profiles

To assess the baseline genetic variation of identical strains when these strains were sequenced in different sequencing institutes, we compared the cgMLST and wgMLST profiles among the strains from the three participating institutes. Only for *E. coli* and *K. pneumoniae* cgMLST schemes were available for use in BioNumerics and, on average, 2441 (97.1%) and 615 (97.1%%) core genome alleles were called for *E. coli* and *K. pneumoniae* respectively (Supplemental Figure 1). In total, 27 and 15 pairwise allelic distances were calculated among the nine *E. coli* and five *K. pneumoniae* strains. In 25/27 (93%) and 12/15 (80%) comparisons, a perfect concordance of cgMLST profiles was observed. In the discordant comparisons, only one allele was differently called between two datasets (Supplemental figure 2).

### Inter-laboratory variation in wgMLST profiles

A total number of 90 pairwise comparisons were made for *K. oytoca* (6), *Citrobacter sp.* (12), *E. coli* (27), *K. pneumoniae* (15), *E. cloacae* (12), *K. aerogenes* (6), *E. faecalis* (6) and *E. faecium* (6). Perfect concordance in wgMLST profiles was obtained in 26/90 (29%) comparisons (Figure 3). In 44/90 (49%) pairwise comparisons, one or two discrepant alleles were observed. Only 23/90 (22%) comparisons showed more than two alleles discrepant, with a maximum of seven alleles different for an *E. coli*. For *E. faecium-1*, which contained the contamination in the dataset, a perfect concordance of wgMLST profiles was observed (data not shown), indicating the robustness of allele-based typing despite contamination with bacterial DNA from different species. For all species, an average allelic distance of 1.6 alleles (standard deviation 1.6) was observed.

**Figure 3.**
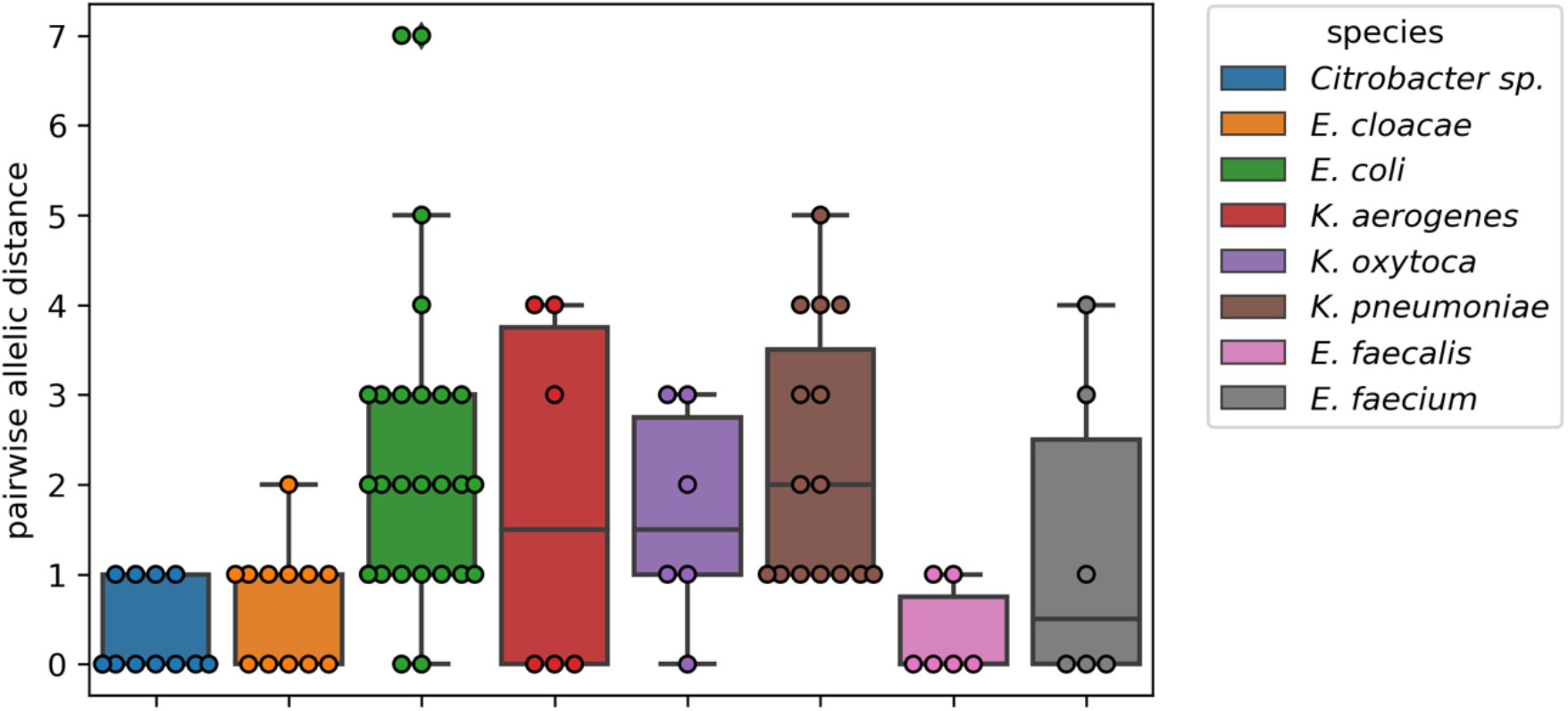
Boxplot of the allelic distance based on wgMLST between the triplicates that were selected for WGS. Boxes show interquartile (IQ) range and whiskers range up to 1.5 times the IQ range. All single pairwise observations were plotted as dots.

For *Citrobacter sp*. a highly diverse number of wgMLST alleles called was observed for the four different strains that were selected for this study. This ranged from 1933 to 4426 alleles that were called, even though the genome size did not vary strongly (mean 4.88Mb, range 4.66Mb to 5.31Mb) and much lower normalized allele counts (mean 0.61, range 0.41 to 0.83) were observed than in other species in this study (mean 0.84, range 0.43 to 0.98). Therefore, determination of the variation on wgMLST for Citrobacter cannot be determined in this study as an incomplete set of alleles were called.

### Reference free wgSNP

As mutations in the genome can also arise in intergenic regions (which are not taken into account in MLST based methods), all assemblies of each isolate were screened using pairwise SNPs. Pairwise SNPs were determined using split k-mers with a variable middle base-pair, as implemented by SKA^13^. First, the most optimal assembler for this task was chosen. For this, we determined the inter- and intra-assembler variation introduced on the number of pairwise SNP between two *de novo* assemblies. The best assembler was chosen based on the one that introduced the least number of pairwise SNPs in the datasets from the same strains with the intra-assembler comparison. To reduce complexity, only the *E. coli* dataset was used. Secondly, the number of pairwise SNPs was determined of the entire dataset employing the best suited assembler. Additionally, also the assembly free method for determining SNPs was used as in implemented by SKA. The mean intra-assembly variation ranged from, 0.2 SNPs (assembly free), 2.7 SNPs (SKESA), 26.6 SNPs (SPAdes), up to 77.8 SNPs (Megahit) (Figure 4ABCD). The mean Inter-assembler variation ranged from 3.9 (assembly free compared to SPAdes) up to 43.0 SNPs (“SPAdes to megahit”). All combinations, except the “assembly-free to assembly-free” and “SKESA to SKESA”, revealed pairwise comparisons with over 20 SNPs for the *E. coli* dataset. Therefore, only these two methods were used to analyze the complete dataset.

**Figure 4.**
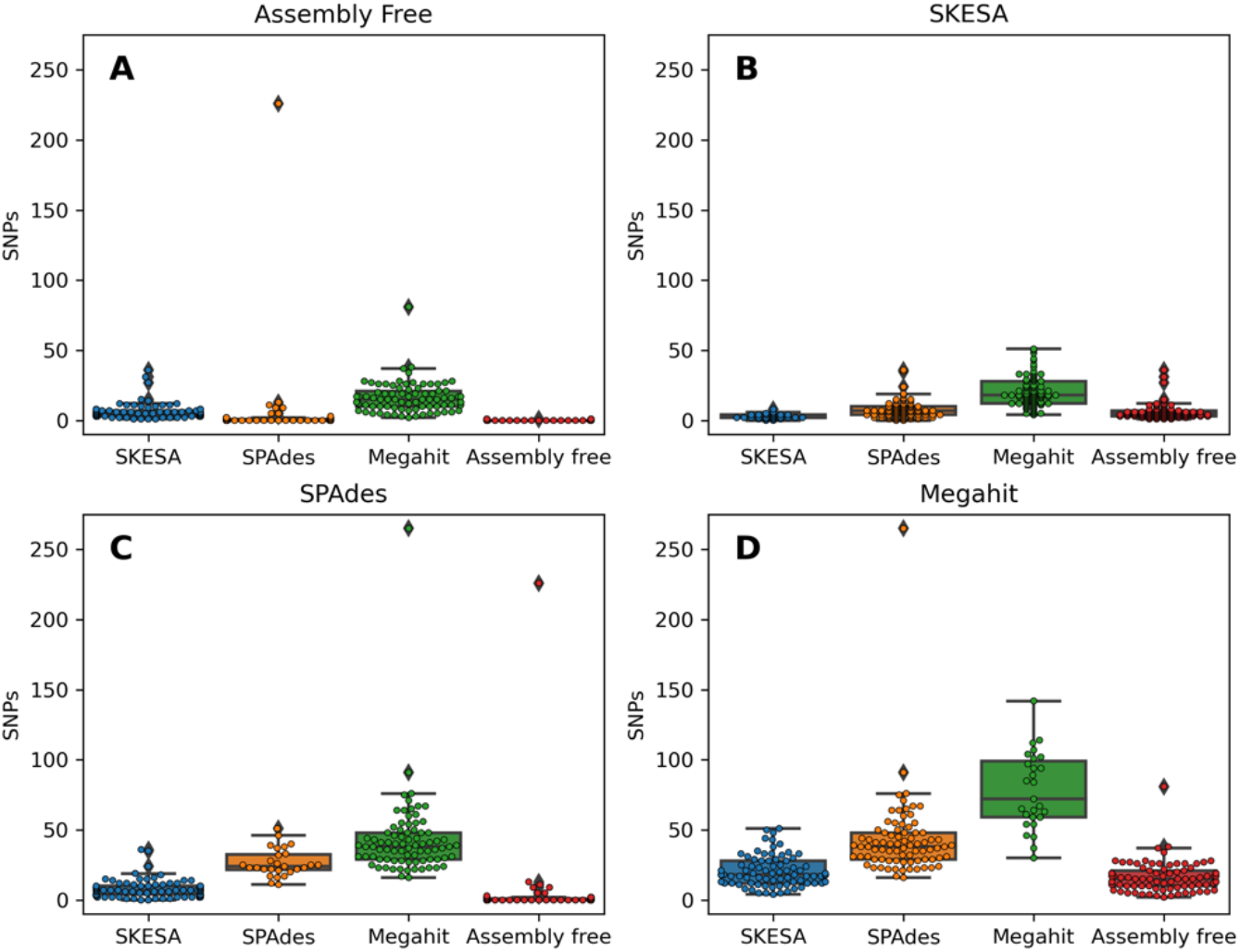
Boxplots of the inter and intra assembly difference in de novo assemblies based on SNPs, using SKA for the *E. coli* dataset. De novo assembly method compared to is indicated above each box. A. Assembly free, B SKESA, C SPAdes and D Megahit. Boxes show interquartile (IQ) range. Whiskers range up to 1.5 times the IQ range. All single pairwise observations were plotted as dots.

Using the assembly free approach, 63/90 (70%) and 21/90 (21%) comparisons show zero or one pairwise SNPs respectively (Figure 5A). Only for *K. pneumoniae*, *E. faecium, K. oxytoca*, and *K. aerogenes* more than 1 pairwise SNP was observed, with a maximum of 5 SNPs for *K. oxytoca*. Using the assembly-based approach, in 10/90 (10%) comparisons, zero SNPs were observed among assemblies (Figure 5B). In 72/90 (80%) of the comparisons less than 5 pairwise SNPs were observed. Interestingly, in the *E. aerogenes* and *K. oxytoca* datasets, more than 8 pairwise SNPs were observed. However, on wgMLST no more than 4 alleles difference was observed. On average 3.4 (± 2.6) pairwise SNPs were observed between assemblies of the same isolates (but sequenced in different institutes). Overall, more pairwise SNPs were observed when assemblies were used for SNP analysis compared to screening raw reads for SNPs. Comparing the number of k-mers between the assembly free and assembly-based methods ranged from −2.1% to 1.2% (Supplemental figure 3), indicating that a similar amount of k-mers between the two methods were compared.

**Figure 5.**
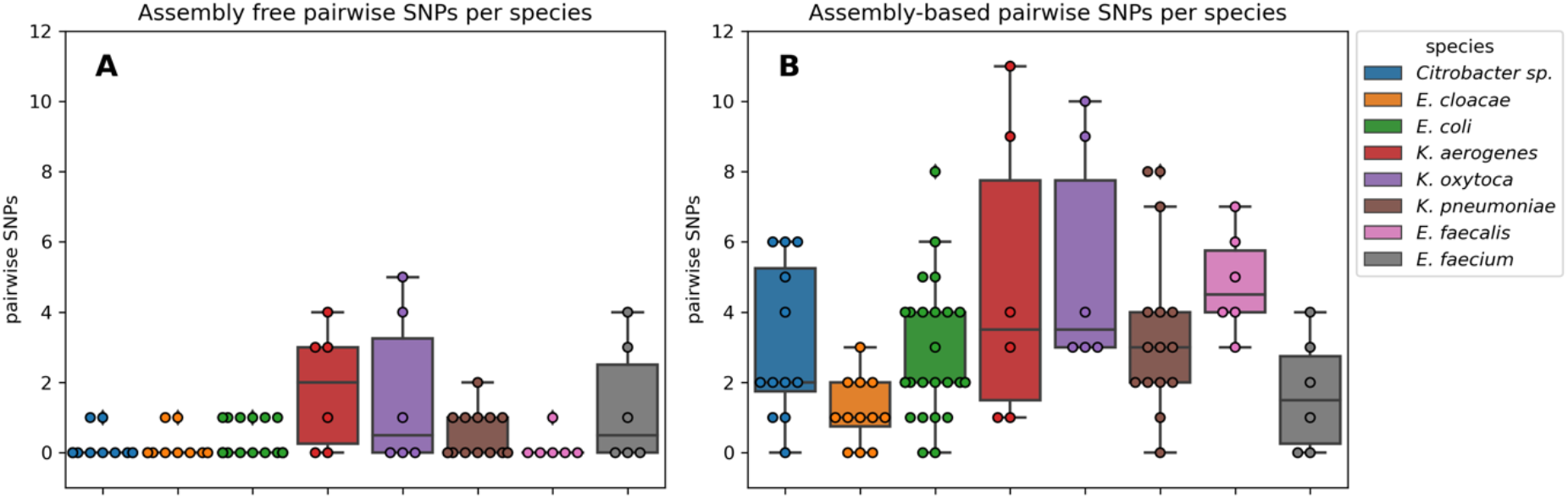
Boxplot of the SNP distance between the triplicates that were selected for WGS. Boxes show interquartile (IQ) range and whiskers range up to 1.5 times the IQ range. All single pairwise observations were plotted as dots. Panel A shows SNP distances using the raw reads as input for SKA. Panel B shows the SNP distances based on the *de novo* assembly using SKESA.

## Discussion

The reproducibility of whole genome sequencing for outbreak surveillance purposes by an inter-laboratory ring-trial was evaluated. Participating institutes subjected the same set of 30 bacterial isolates of various Enterobacteriaceae and Enterococci species for whole genome sequencing. As a first step, we assessed various QC measures. We observed a slight positive trend of the sequencing depth on the normalized number of alleles called in the sequencing depth range of 30 to 207-fold. It remains unclear what a sufficient sequencing depth is to correctly reconstruct the maximum number of correct alleles in the genome. Kluytmans van den Bergh et al.^21^ demonstrated an increase in resolution for phylogenetic reconstruction of Enterobacteriaceae if wgMLST is implemented compared to cgMLST. This would indicate that more alleles available for comparison will improve the surveillance of outbreaks by cgMLST or wgMLST methods. Therefore, it is advisable to generate WGS data of sufficient depth to maximize the number of loci in the *de novo* assembly. On the other hand, deeper sequencing after a certain depth, may not improve the phylogenetic signal any further, and does increase run-time of subsequent *de novo* assembly.

Additionally, generally we observed a lower than expected allele density than expected. Prokaryotes show a coding density of 1 CDS per 1 kb^35^. The majority of datasets show a lower number allele density (0.83 per kb, Figure 1D), which could be caused by the quality filtering step in allele calling. Yet, the low number of called alleles for most *Citrobacter sp.* may be explained by incomplete allele schemes, which do not contain the complete diversity of alleles. This indicates the low representation of the complete diversity of *Citrobacter sp.* genome assemblies present in public databases and may indicate the finding of new antibiotic-resistant *Citrobacter sp*. in The Netherlands.

### Genotyping AMR genes and ORIs

Next, Identification of AMR genes and plasmid ORIs was performed. Overall, an excellent reproducibility was achieved, as a precision of 99% was obtained. Most discrepancies could be explained by the variation in the variant calling of a specific resistance gene. Four times in 973 genotype calls, there was an unexplained absence of a resistance gene. Although the DNA isolation method used here showed good results for the application of WGS^36^, still some loci could be missed due to inefficient isolation of plasmid DNA where these AMR genes can be located. Only twice and in different institutes, an unexpected ColpVC ORI was found in one of the sequencing datasets, which may indicate contamination during DNA isolation or library preparation. Strauß and co-authors reported a 1.7% discordance caused by WGS compared to micro-array for the detection of resistance and virulence genes^37^. In this study, a similar reproducibility in typing resistance genes and ORIs was obtained and previously described by Kozyreva *et al.,* which found a reproducibility rate of 99.97% ^38^.

### Genetic variation

It is of great importance that the genetic distance between technical duplicates does not surpass commonly used thresholds to classify strains into clusters. Here, some variation among the wgMLST allelic profiles was observed, which translated to an average of 0.49 discrepant alleles per 1000 alleles compared. Kluytmans-Van den Bergh *et al*.^21^ reported a variation in genetic distance based on wgMLST in a range of 0 to 0.001 (which translates to 5 alleles difference, based on 5000 alleles compared) for five *E. coli* and three *K. pneumoniae* which were sequenced in duplicate^21^. This is in concordance with our study, where 88/90 comparisons have no more than 5 alleles difference. Additionally, reported clonal thresholds by these authors were roughly 26 and 2 alleles difference for *E. coli* and *K. pneumoniae* on cgMLST respectively. For wgMLST this was 29, 23, 8, 14 alleles difference for *E. coli, K. pneumoniae*, *Citrobacter sp.* and *Enterobacter cloacae* complex respectively. These clonal thresholds are higher by a safe margin than the variation between any of the replicates, sequenced of datasets. Although variation on a genetic level was observed, the level of disparity remained below other thresholds commonly employed for hospital outbreak surveillance purposes ^9^. Previous work suggested a cut-off of 10 alleles for MDR *E. coli* and *K. pneumoniae* based on cgMLST^39,40^. Therefore, it is safe to assume that if harmonized protocols are used, the genetic difference remains within these previously described thresholds.

In the wgSNP analysis, all methods except for the “assembly-free to assembly-free” and “SKESA to SKESA” showed pairwise comparisons with more than 20 SNPs. This indicates that using SPAdes or Megahit in combination with a SNP based method, is unsuitable for outbreak surveillance, as identical strains have more SNPs than commonly used outbreak thresholds^9^, indicating that these strains would be considered not clonally related, thus not belonging to the same outbreak. Furthermore, this also held true when comparing two assembly methods, which implies that comparing bacterial assemblies should be avoided at all costs, if multiple centers employ different methodologies to generate *de novo* assemblies for WGS outbreak surveillance. Potential outbreak can be missed due to the large number of SNPs detected results in identical isolates not being flagged as clonally related, which subsequently have implications for infection prevention control. For the assembly free method, we observed most replicates to have no SNPs between each other (70%), which is in line with the GenomeTrakr proficiency-test study, which found a similar fraction of datasets showed having no SNPs (73%)^41^.

Variation in SNPs among strains showed lower number of SNPs based on the assembly-free method compared to the assembly-based method. It is unlikely that this is caused by different number of k-mers that were compared for SNPs, as there was only a modest difference for the number of k-mers compared between the assembly free and the assembly based SNP analysis, which ranged from - 2.1% to 1.2% difference in compared k-mers (Supplemental figure 3). Therefore, it is more likely that *de novo* assembly introduces phylogenetic noise in regions difficult to assemble, i.e. around regions such as mobile elements (*i.e.* transposons and plasmids). Previously described work employing SNP-based methodologies to infer phylogeny among bacterial isolates often mask regions in the genome that are sensitive for non-informative SNPs for phylogenetic reconstruction, such as mobile genetic elements (MGE). Masking of these regions requires specialized tools such as Gubbins^42^ that are able to recognize regions with elevated numbers of base substitutions in the genome. Unfortunately, using this reference-free methodology, makes this masking impossible to perform in an unbiased and automated fashion, like in the Gubbins pipeline. Therefore, we must assume the possibility of overestimation of SNPs among strains in our study.

### Study limitations

For this study, only three centers participated in this ring-trial, all part of the I-4-1-Health study group. Here, ESBL-producing and ciprofloxacin-resistant Enterobacteriaceae and vancomycin-resistant *Enterococcus* were defined of primordial interest, however other important nosocomial bacterial pathogens such as *Pseudomonas* sp., *Staphylococci* and A*cinetobacter sp.* were not included in the study. Furthermore, all three centers used the same protocols for the extraction and library preparation for sequencing on the Illumina MiSeq. Recommendations for future research would therefore be to determine if these harmonized wetlab protocols and subsequent bioinformatic data processing are indeed required for the reconstruction of outbreak clusters.

### Conclusion

Overall, the work presented here demonstrated that whole genome sequencing generates reproducible results when comparing results from an inter-laboratory perspective using identical wet-lab and dry-lab methodologies for WGS. Furthermore, to make multi-center outbreak surveillance feasible in the future, we recommend to share raw sequencing reads as systematic errors were introduced in the *de novo* assemblies among different assemblers. Finally, work presented here lays the foundation for routine proficiency testing in clinical microbiology laboratories.

## Acknowledgements

We are grateful to the collaborators in the participating laboratories for their contribution to the collection of whole genome sequence data.

## Supplemental figures

**Supplemental figure 1.**
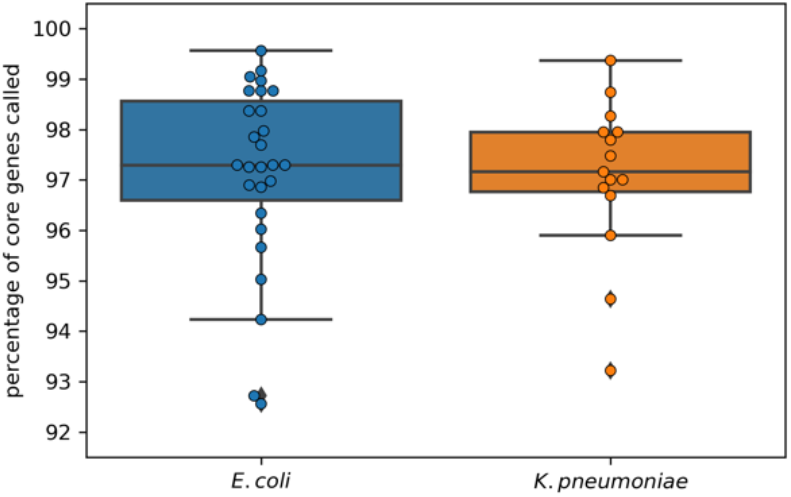
Boxplot of the number of core genes called for cgMLST in percentage for *K. pneumoniae* and *E. coli* respectively. Boxes range the interquartile (IQ) range. Whiskers range up to 1.5 times the IQ range. All single datapoints are represented as single dots. Only cgMLST schemes were available for *E. coli* and *K. pneumoniae*.

**Supplemental figure 2.**
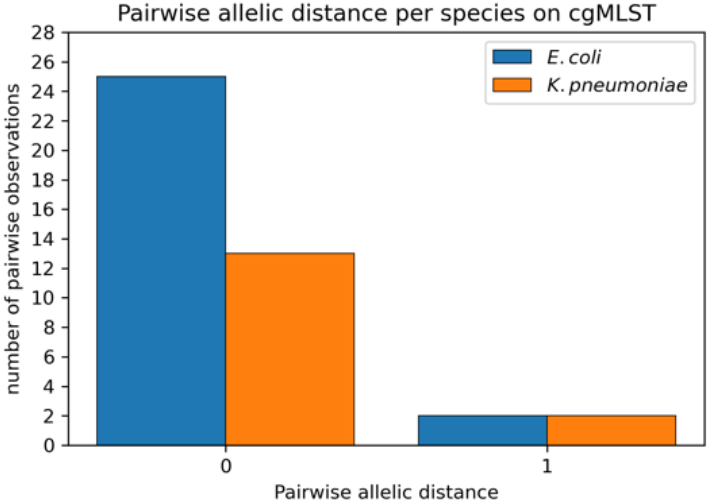
Barplot of the pairwise number of alleles that were different between two strains.

**Supplemental figure 3.**
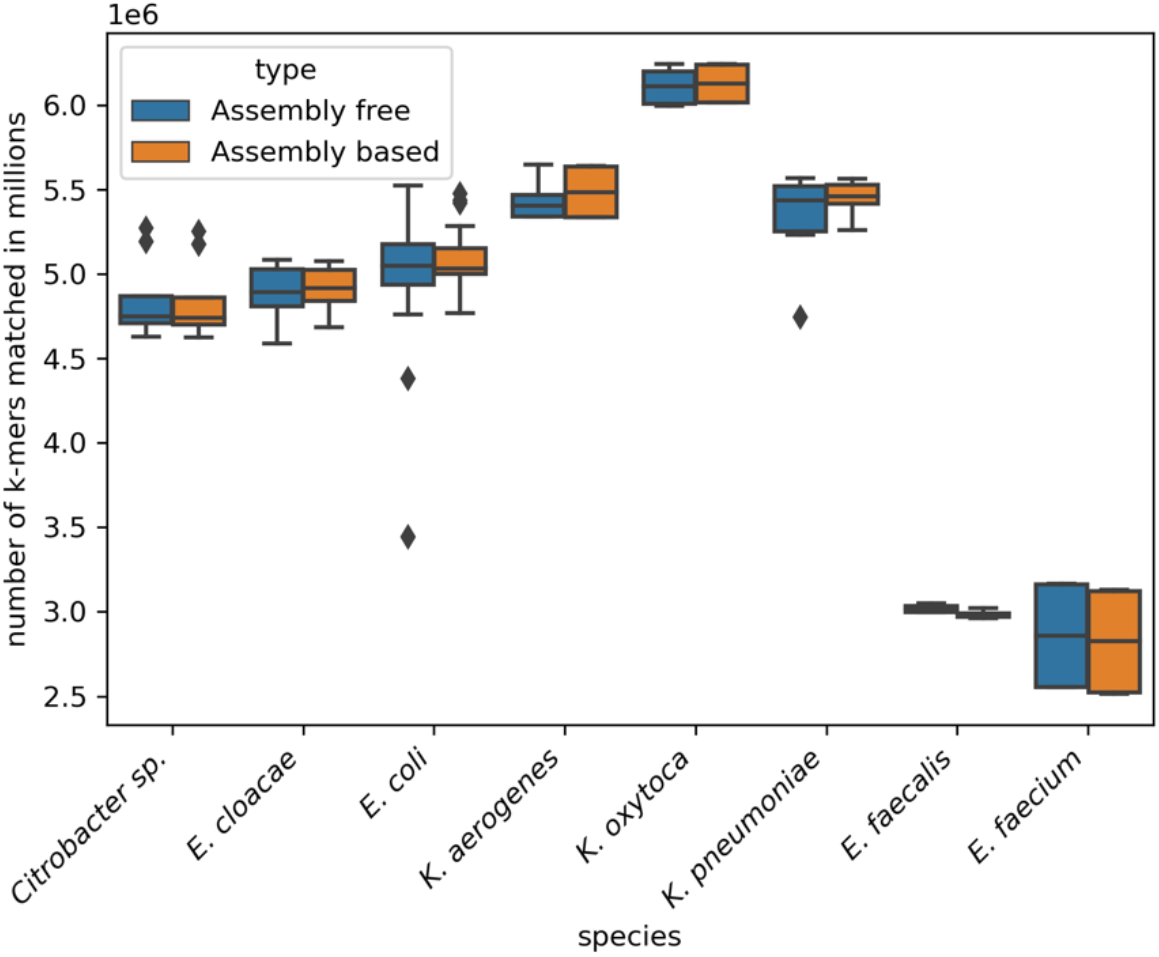
boxplot of the number of k-mers compared for the assembly free and the assembly based method. Boxes range the interquartile (IQ) range. Whiskers range up to 1.5 times the IQ range. Outliers were shown as single datapoints.

## References

1. Global action plan on antimicrobial resistance. https://www.who.int/publications-detail/global-action-plan-on-antimicrobial-resistance. Accessed March 13, 2020.

2. Thaden JT, Li Y, Ruffin F, et al. Increased costs associated with bloodstream infections caused by multidrug-resistant gram-negative bacteria are due primarily to patients with hospital-acquired infections. Antimicrob Agents Chemother. 2017. doi:10.1128/AAC.01709-16

3. Founou RC, Founou LL, Essack SY. Clinical and economic impact of antibiotic resistance in developing countries: A systematic review and meta-analysis. PLoS One. 2017. doi:10.1371/journal.pone.0189621

4. Mariappan S, Sekar U, Kamalanathan A. Carbapenemase-producing Enterobacteriaceae: Risk factors for infection and impact of resistance on outcomes. Int J Appl Basic Med Res. 2017. doi:10.4103/2229-516x.198520

5. Harris SR, Feil EJ, Holden MTG, et al. Evolution of MRSA during hospital transmission and intercontinental spread. Science (80-). 2010. doi:10.1126/science.1182395

6. Pendleton S, Hanning I, Biswas D, Ricke SC. Evaluation of whole-genome sequencing as a genotyping tool for Campylobacter jejuni in comparison with pulsed-field gel electrophoresis and flaA typing. Poult Sci. 2013. doi:10.3382/ps.2012-02695

7. Leekitcharoenphon P, Nielsen EM, Kaas RS, Lund O, Aarestrup FM. Evaluation of whole genome sequencing for outbreak detection of salmonella enterica. PLoS One. 2014. doi:10.1371/journal.pone.0087991

8. Salipante SJ, SenGupta DJ, Cummings LA, Land TA, Hoogestraat DR, Cookson BT. Application of whole-genome sequencing for bacterial strain typing in molecular epidemiology. J Clin Microbiol. 2015. doi:10.1128/JCM.03385-14

9. Schürch AC, Arredondo-Alonso S, Willems RJL, Goering R V. Whole genome sequencing options for bacterial strain typing and epidemiologic analysis based on single nucleotide polymorphism versus gene-by-gene–based approaches. Clin Microbiol Infect. 2018. doi:10.1016/j.cmi.2017.12.016

10. Silva M, Machado MP, Silva DN, et al. chewBBACA: A complete suite for gene-by-gene schema creation and strain identification. Microb genomics. 2018. doi:10.1099/mgen.0.000166

11. Seemann T. Snippy. https://github.com/tseemann/snippy.

12. Wu TD, Nacu S. Fast and SNP-tolerant detection of complex variants and splicing in short reads. Bioinformatics. 2010;26. doi:10.1093/bioinformatics/btq057

13. Harris SR. SKA: Split Kmer Analysis Toolkit for Bacterial Genomic Epidemiology. bioRxiv. 2018. doi:10.1101/453142

14. Hyatt D, Chen GL, LoCascio PF, Land ML, Larimer FW, Hauser LJ. Prodigal: Prokaryotic gene recognition and translation initiation site identification. BMC Bioinformatics. 2010. doi:10.1186/1471-2105-11-119

15. Zhou Z, Alikhan NF, Mohamed K, Fan Y, Achtman M. The EnteroBase user’s guide, with case studies on Salmonella transmissions, Yersinia pestis phylogeny, and Escherichia core genomic diversity. Genome Res. 2020. doi:10.1101/gr.251678.119

16. Darling AE, Mau B, Perna NT. Progressivemauve: Multiple genome alignment with gene gain, loss and rearrangement. PLoS One. 2010. doi:10.1371/journal.pone.0011147

17. Treangen TJ, Ondov BD, Koren S, Phillippy AM. The harvest suite for rapid core-genome alignment and visualization of thousands of intraspecific microbial genomes. Genome Biol. 2014. doi:10.1186/s13059-014-0524-x

18. Gardner SN, Slezak T, Hall BG. kSNP3.0: SNP detection and phylogenetic analysis of genomes without genome alignment or reference genome. Bioinformatics. 2015. doi:10.1093/bioinformatics/btv271

19. Harris SR, Cartwright EJP, Török ME, et al. Whole-genome sequencing for analysis of an outbreak of meticillin-resistant Staphylococcus aureus: A descriptive study. Lancet Infect Dis. 2013. doi:10.1016/S1473-3099(12)70268-2

20. Rumore J, Tschetter L, Kearney A, et al. Evaluation of whole-genome sequencing for outbreak detection of Verotoxigenic Escherichia coli O157:H7 from the Canadian perspective. BMC Genomics. 2018. doi:10.1186/s12864-018-5243-3

21. Kluytmans-Van Den Bergh MFQ, Rossen JWA, Bruijning-Verhagen PCJ, et al. Whole-genome multilocus sequence typing of extended-spectrum-beta-lactamase-producing enterobacteriaceae. J Clin Microbiol. 2016. doi:10.1128/JCM.01648-16

22. Bergh MK Den, Lammens C, Selva NP, et al. Microbiological methods to detect intestinal carriage of highly-resistant microorganisms (HRMO) in humans and livestock in the i-4-1-Health Dutch-Belgian cross-border project. Preprints.org. 2019;(December):1–16. doi:10.20944/preprints201912.0216.v1

23. Bergh MK Den, Lammens C, Selva NP, et al. Microbiological methods to detect intestinal carriage of highly-resistant microorganisms (HRMO) in humans and livestock in the i-4-1-Health Dutch-Belgian cross-border project. Preprints.org. 2019;(December). doi:10.20944/preprints201912.0216.v1

24. Virtanen P, Gommers R, Oliphant TE, et al. SciPy 1.0: fundamental algorithms for scientific computing in Python. Nat Methods. 2020. doi:10.1038/s41592-019-0686-2

25. Bankevich A, Nurk S, Antipov D, et al. SPAdes: A new genome assembly algorithm and its applications to single-cell sequencing. J Comput Biol. 2012. doi:10.1089/cmb.2012.0021

26. Altschul SF, Gish W, Miller W, Myers EW, Lipman DJ. Basic local alignment search tool. J Mol Biol. 1990. doi:10.1016/S0022-2836(05)80360-2

27. Zankari E, Hasman H, Cosentino S, et al. Identification of acquired antimicrobial resistance genes. J Antimicrob Chemother. 2012. doi:10.1093/jac/dks261

28. Carattoli A, Zankari E, Garciá-Fernández A, et al. In Silico detection and typing of plasmids using plasmidfinder and plasmid multilocus sequence typing. Antimicrob Agents Chemother. 2014. doi:10.1128/AAC.02412-14

29. Souvorov A, Agarwala R, Lipman DJ. SKESA: Strategic k-mer extension for scrupulous assemblies. Genome Biol. 2018. doi:10.1186/s13059-018-1540-z

30. Li D, Liu CM, Luo R, Sadakane K, Lam TW. MEGAHIT: An ultra-fast single-node solution for large and complex metagenomics assembly via succinct de Bruijn graph. Bioinformatics. 2015. doi:10.1093/bioinformatics/btv033

31. Köster J, Rahmann S. Snakemake-a scalable bioinformatics workflow engine. Bioinformatics. 2012. doi:10.1093/bioinformatics/bts480

32. Tae H, Karunasena E, Bavarva JH, McIver LJ, Garner HR. Large scale comparison of non-human sequences in human sequencing data. Genomics. 2014;104(6):453–458. doi:10.1016/j.ygeno.2014.08.009

33. Sangiovanni M, Granata I, Thind AS, Guarracino MR. From trash to treasure: Detecting unexpected contamination in unmapped NGS data. BMC Bioinformatics. 2019. doi:10.1186/s12859-019-2684-x

34. Glassing A, Dowd SE, Galandiuk S, Davis B, Chiodini RJ. Inherent bacterial DNA contamination of extraction and sequencing reagents may affect interpretation of microbiota in low bacterial biomass samples. Gut Pathog. 2016. doi:10.1186/s13099-016-0103-7

35. Koonin E V., Wolf YI. Genomics of bacteria and archaea: The emerging dynamic view of the prokaryotic world. Nucleic Acids Res. 2008;36(21):6688–6719. doi:10.1093/nar/gkn668

36. Nouws S, Bogaerts B, Verhaegen B, et al. Impact of DNA extraction on whole genome sequencing analysis for characterization and relatedness of Shiga toxin-producing Escherichia coli isolates. Sci Rep. 2020. doi:10.1038/s41598-020-71207-3

37. Strauß L, Ruffing U, Abdull S, et al. Detecting staphylococcus aureus virulence and resistance genes: A comparison of whole-genome sequencing and DNA microarray technology. J Clin Microbiol. 2016. doi:10.1128/JCM.03022-15

38. Kozyreva VK, Truong CL, Greninger AL, Crandall J, Mukhopadhyay R, Chaturvedi V. Validation and implementation of clinical laboratory improvements act-compliant whole-genome sequencing in the public health microbiology laboratory. J Clin Microbiol. 2017. doi:10.1128/JCM.00361-17

39. Roer L, Hansen F, Frølund Thomsen MC, et al. WGS-based surveillance of third-generation cephalosporin-resistant Escherichia coli from bloodstream infections in Denmark. J Antimicrob Chemother. 2017. doi:10.1093/jac/dkx092

40. Zhou H, Liu W, Qin T, Liu C, Ren H. Defining and evaluating a core genome multilocus sequence typing scheme for whole-genome sequence-based typing of Klebsiella pneumoniae. Front Microbiol. 2017. doi:10.3389/fmicb.2017.00371

41. Timme RE, Rand H, Leon MS, et al. GenomeTrakr proficiency testing for foodborne pathogen surveillance: An exercise from 2015. Microb Genomics. 2018. doi:10.1099/mgen.0.000185

42. Croucher NJ, Page AJ, Connor TR, et al. Rapid phylogenetic analysis of large samples of recombinant bacterial whole genome sequences using Gubbins. Nucleic Acids Res. 2015. doi:10.1093/nar/gku1196

